# Prowler: A novel trimming algorithm for Oxford Nanopore sequence data

**DOI:** 10.1101/2021.05.09.443332

**Authors:** Simon Lee, Loan T Nguyen, Ben J Hayes, Elizabeth Ross

## Abstract

**Motivation:** Quality control (QC) tools are critical in DNA sequencing analysis because they increase the accuracy of sequence alignments and thus the reliability of results. Oxford Nanopore Technologies (ONT) QC is currently rudimentary, generally based on whole read average quality. This results in discarding reads that contain regions of high quality sequence. Here we propose Prowler, a multi-window approach inspired by algorithms used to QC short read data. Importantly, we retain the phase and read length information by optionally replacing trimmed sections with Ns.

**Results:** Prowler was applied to mammalian and bacterial datasets, to assess effects on alignment and assembly respectively. Compared to Nanofilt, alignments of data QC’ed with Prowler had lower error rates and more mapped reads. Assemblies of Prowler QC’ed data had a lower error rate than Nanofilt QCed data however this came at some cost to assembly contiguity.

**Availability and implementation:** Prowler is implemented in Python and is available at: https://github.com/ProwlerForNanopore/ProwlerTrimmer

**Supplementary information:** Supplementary data are available at Bioinformatics online.

## 1 Introduction

Quality control (QC) tools are critical in DNA sequencing because they increase the accuracy of sequence alignments and thus the reliability of results. The current best practice for performing QC on Oxford Nanopore Technologies (ONT) data is to take the average Phred Quality Score of the entire read and either accept or reject based on an average quality score cut-off (Rang et al., 2018, Loman & Quinlan, 2014), implemented in Nanofilt (De Coster *et al*., 2018). For ONT reads that vary in quality, an average Q-Score discards reads even with long stretches of high quality sequence. An algorithm that is able to salvage usable sequences would make best use of the current generation of sequencing technologies.

Here we present an alternative trimming tool that can reduce the error rate of alignment and assembly of ONT by dividing the sequence into windows and assessing the quality of each window: Prowler.

## 2 Methods

### Prowler Design

Prowler (PROgressive multi-Window Long Read trimmer) was developed to remove low average Q-Score segments. The Prowler algorithm (Figure 1A) considers the quality distribution of the read by breaking the sequence into multiple non-overlapping windows.

**Figure 1:**
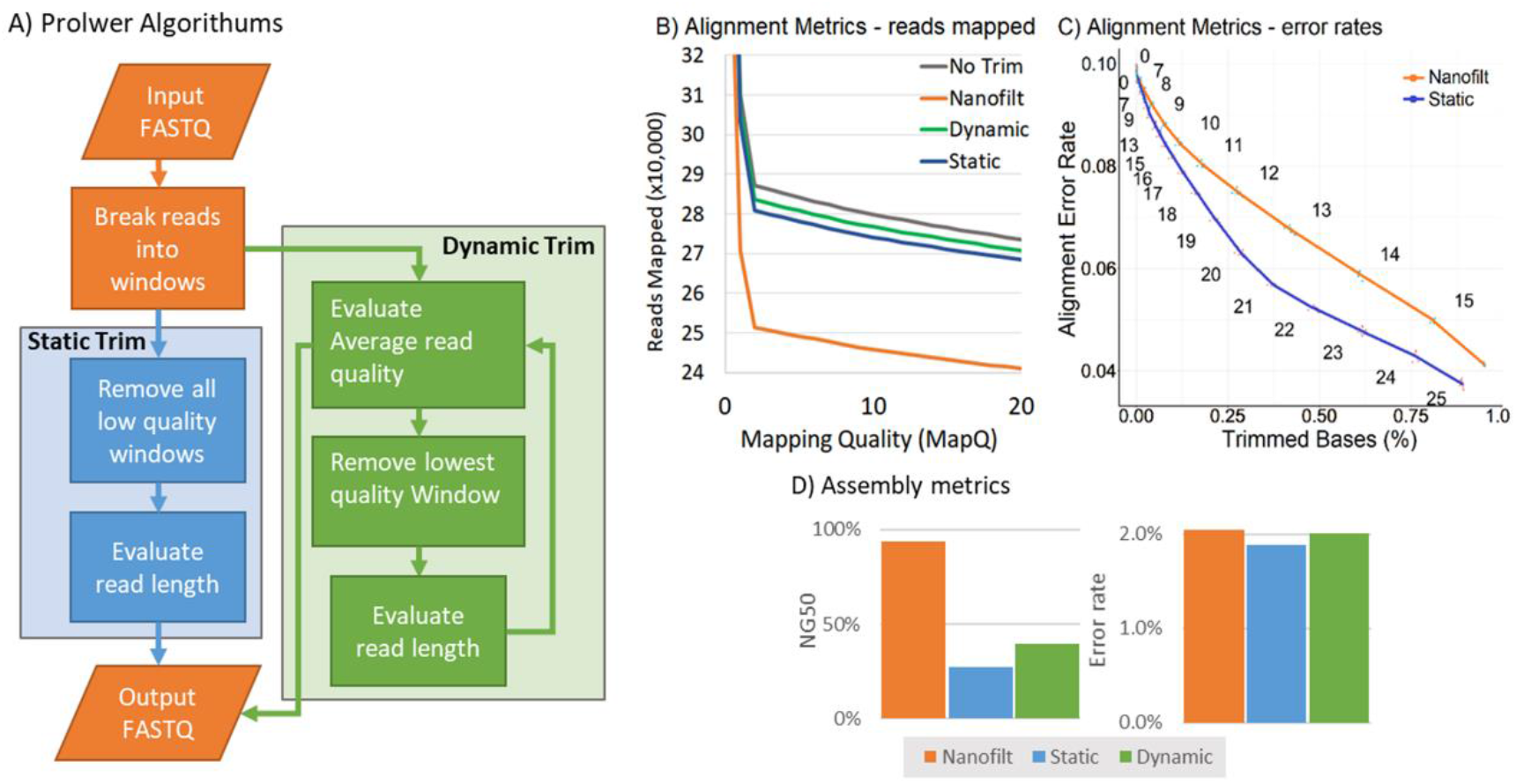
A) Prowler algorithm. B) Mapping rates after quality control of Bos indicus data. C) Error rate of mapped reads in Bos indicus data. D) S. syrphidae assembly metrics – NG50 shows as % of full genome length.

Prowler was designed with two algorithms: static and dynamic. In the static algorithm, Prowler removes all windows that fall below a given threshold. In the dynamic algorithm, Prowler progressively removes the lowest quality window before recalculating average whole-read quality. When average whole-read quality passes a minimum threshold, the sequence is accepted. In both static and dynamic algorithm mode, if the total number of remaining bases falls below a minimum number, the read is discarded. Dynamic Prowler salvages reads that would be rejected by Nanofilt, while Static Prowler also improves the quality of previously accepted reads by applying a fixed trim at the window level.

Prowler has two output modes: U0 outputs unfragmented reads - low quality regions are output as a string of N of equal length to the replaced window; FX outputs the longest X continuous fragments that meet the trimming criteria (e.g. F1 outputs one fragment per read, F2 outputs up to two fragments per read, and so forth).

### Algorithm testing

The effects of trimming ONT data with Prowler were tested on two datasets: 10GB of *Bos indicus* (cattle) MinION data (9.4.1 flowcell; guppy v4.1 basecaller; (Ross, 2019)), and 400MB of *S. syrphidae* MinION data (NZ_CP025257.1; SAMN08158127, SRR6364637_1;(Knight, 2017)).

Data was filtered using Nanofilt and Prowler both with minimum length of 1000bp. The alignment quality was compared by aligning the data using Minimap2 (Li, 2018), with –score-N 0. Bacterial data was assembled with Canu (Koren *et al*., 2017). See Supplementary document for Prowler settings. Nanofilt was run with TrimQ from 0-16.

## 3 Results

When tested with 4 CPUs and 4GB of RAM Prowler processed 16000 reads per minute. Dynamic Prowler was 10% faster than Static Prowler. With a quality threshold of 10, on the *B. indicus* data with a window size of 500 and a minimum read length of 1000, Nanofilt discarded 16.7% of reads, while U0-Prowler static discarded 7.2% and U0-Prowler dynamic discarded 5.9%. When the reads were mapped to the reference genome more reads mapped with high accuracy (MapQ>20) after QC with Prolwer than with Nanofilt (Figure 1B). The error rate of the alignments was lower after QC with Prowler compared to Nanofilt (Figure 1C).

When *S. syrphidae* data was assembled with Canu after trimming with Prowler and Nanofilt, genomes with > 95% length were assembled. Prowler maintained a high NG50 (>95% of genome length) while achieving the lowest error rate (2.14%). Static F1 Prowler achieved the best error reduction (P < 0.001) at the cost of NG50 falling to 28.2% of reference genome size (Figure 1D). Using Prowler beyond a certain TrimQ causes the reads to become too fragmented, resulting in a decrease in NG50.

## 4 Conclusion

Here we present Prowler, a QC program enabling recovery of high quality windows of otherwise low quality Nanopore reads. Prowler out-performs Nanofilt as a QC program for ONT reads. The specific settings that are applied need to be considered when selecting trimming settings for Prowler due to the tradeoff between continuality and error rate of assemblies. Prowler is able to increase the data yield of ONT devices by salvaging high quality regions of reads that would otherwise be discarded based on the entire read average quality.

## Supporting information

Prowler assemble and alignment data

## Acknowledgements

The Authors gratefully acknowledge statistical advice from Dr Roy Costilla.

